# A second PI(4,5)P_2_ binding site determines PI(4,5)P_2_ sensitivity of the tubby domain

**DOI:** 10.1101/2020.09.23.309492

**Authors:** Veronika Thallmair, Lea Schultz, Siewert J. Marrink, Dominik Oliver, Sebastian Thallmair

**Affiliations:** Department of Neurophysiology, Institute of Physiology and Pathophysiology, Philipps-University Marburg, Deutschhausstr. 1-2, 35037 Marburg, Germany; DFG Research Training Group, Membrane Plasticity in Tissue Development and Remodeling, GRK 2213, Philipps University, Germany; Groningen Biomolecular Sciences and Biotechnology Institute and Zernike Institute for Advanced Materials, University of Groningen, Nijenborgh 7, 9747 AG Groningen, Netherlands

**Keywords:** tubby, phosphoinositide, PIP2, TULP, membrane interface, cilium, Martini model

## Abstract

Phosphosinositides (PIs) are lipid signaling molecules that operate by recruiting proteins to cellular membranes via PI recognition domains. Such domains are also used widely as fluorescence-coupled biosensors for cellular PIs. For PI(4,5)P_2_, the dominant PI of the plasma membrane (PM), only two recognition domains have been characterized in detail and used as sensors. One of them, the tubby domain, which is conserved in the tubby-like protein (TULP) family, is essential for targeting proteins into cilia in a process involving reversible membrane association. However, the PI(4,5)P_2_ binding properties of tubby domains have remained enigmatic.

Here we used coarse-grained molecular dynamics (MD) simulations to explore PI(4,5)P_2_ binding by the prototypic tubby domain (tubbyCT). While the MD simulations showed a comparatively low PI(4,5)P_2_ affinity of the previously described canonical binding site, they unexpectedly revealed an adjacent second binding site, consisting of a conserved cationic cluster at the protein-membrane interface. Population of this second site dramatically increased membrane association of tubbyCT. Although less specific than the canonical binding pocket, this second site preferred binding of PI(4,5)P_2_ over PI(4)P and phosphatidyl serine. Mutations in this site impaired PI(4,5)P_2_-dependent PM localization in living cells and PI(4,5)P_2_ interaction *in silico*.

Thus, the second binding site essentially contributes to the effective affinity and hence PM association of the tubby domain. The two-ligand binding mode may serve to sharpen the membrane association-dissociation cycle of TULPs that underlies delivery of ciliary cargo.

## INTRODUCTION

Among the phospholipids, the phosphoinositides (PIs) have multifaceted signaling functions. First, PIs are a fundamental part of the cell’s membrane identity code in eukaryotic cells (Di Paolo and De Camilli, 2006; Dickson and Hille, 2019). Moreover, temporal changes in PI concentrations, in particular at the plasma membrane (PM), instruct important signal transduction pathways. Prominent examples are the generation of PIP3 downstream of growth factor receptors and the depletion of PI(4,5)P_2_ by PLCβ downstream of Gq-coupled receptors. Generally, the impact of PIs on cellular processes is mediated by the binding of proteins to the membrane-localized PI via PI-recognition domains (reviewed by (Balla, 2013; Dickson and Hille, 2019; Hammond and Balla, 2015)). A diversity of such PI-binding domains has been discovered, including PH, PX, FYVE, and ENTH domains among others (Hammond and Balla, 2015; Lemmon, 2008). Importantly, some of these domains bind to a single PI species with high specificity, whereas others are less specific and interact with a range of PIs (McLaughlin and Murray, 2005) or even with anionic lipids other than PIs (Hammond and Balla, 2015; Lemmon, 2008).

Beyond their eminent role in cell biology, ligand-specific PI-binding domains have emerged as highly useful biosensors for their cognate PI lipid in living cells. Encoded genetically to yield fusions with fluorescent (or luminescent) modules such as GFP, they are being used widely to interrogate PI cell biology in model systems across a range of biological complexity from isolated membrane fragments (Milosevic et al., 2005) to intact living organisms/animals (Hardie et al., 2015). The general principle is that binding of such a sensor to a membrane reports on the presence of the recognized lipid at a biologically relevant concentration. Accordingly, translocation of the probe to or from the membrane reports on dynamic changes in the recognized lipid’s concentration (reviewed in (Hammond and Balla, 2015)).

PI(4,5)P_2_ is the most abundant PI of the PM and besides being a precursor of the canonical second messengers PIP_3_, Ins(1,4,5)P_3_ (IP_3_), and diacylglycerol (DAG) has multiple roles as a bon-fide second messenger (Balla, 2013). In fact, the first lipid biosensor invented was a PI(4,5)P_2_-specific PH domain, the PH domain of phospholipase δ1 (PLCδ1-PH) (Stauffer et al., 1998; Várnai and Balla, 1998) that has been used since then in countless studies (reviewed in (Hammond and Balla, 2015)). However, because PLCδ1-PH also binds IP_3_, which can lead to ambiguity in interpretation with respect to PI(4,5)P_2_ dynamics (Hirose et al., 1999), there has been high interest in alternative sensors with better specificity. Surprisingly, only few specific PI(4,5)P_2_ binding domains have been identified (Hammond and Balla, 2015), and fewer have been characterized in depth and have been used as biosensors (Leitner et al., 2019; Szentpetery et al., 2009; Yoon et al., 2011). The only alternative PI(4,5)P_2_ sensor that has been used fairly frequently is the C-terminal domain of the tubby protein (‘tubby domain’, hereafter abbreviated tubbyCT). TubbyCT has been employed to detect PM PI(4,5)P_2_ dynamics in cell culture (Mavrantoni et al., 2015; Quinn et al., 2008), isolated neurons (Nelson et al., 2008), native neurons in brain slices (Hackelberg and Oliver, 2018), and drosophila photoreceptors (Hardie et al., 2015).

A crystal structure of tubbyCT identified a PI(4,5)P_2_ binding site, which is conserved across the tubby domains of tubby-like proteins (TULPs) (Santagata et al., 2001). Notwithstanding, there is considerable confusion concerning the PI(4,5)P_2_ binding properties of tubbyCT, as in some cell types, tubbyCT resists dissociation from the PM during strong activation of PLCβ-mediated depletion of PI(4,5)P_2_, despite confirmation of PI(4,5)P_2_ loss by independent readouts such as PLCδ1-PH (Leitner et al., 2019; Quinn et al., 2008; Szentpetery et al., 2009). This has been taken to indicate a higher PI(4,5)P_2_ affinity of tubbyCT compared to PLCδ1-PH. However, titration of PI(4,5)P_2_ with a voltage-activated PLCδ1-PH phosphatase (Ci-VSP; (Murata et al., 2005)) provided evidence that the affinity of tubbyCT for PI(4,5)P_2_ is actually lower compared to PLCδ1-PH (Halaszovich et al., 2009; Leitner et al., 2019).

This prompted us to employ coarse-grained (CG) molecular dynamics (MD) simulations to explore binding of tubbyCT to PI(4,5)P_2_ in a realistic membrane environment. CG MD simulations are a suitable and well-established tool to study protein-lipid interactions (Corradi et al., 2019). In particular, protein-PI interactions have been successfully modelled using the CG Martini force field (Corradi et al., 2018; Naughton et al., 2016; Naughton et al., 2018; Sun et al., 2020; Yamamoto et al., 2020; Yamamoto et al., 2016). Our simulations of membrane binding showed that tubbyCT’s PI(4,5)P_2_ binding affinity is lower than that of PLCδ1-PH, confirming previous experimental data. Unexpectedly, the MD data revealed a second PI(4,5)P_2_ binding site within tubbyCT’s membrane-oriented surface, comprising a cluster of positively charged residues. Mutation of single amino acids within this second binding site strongly reduced PI(4,5)P_2_ binding of tubbyCT both in-silico and experimentally in living cells, demonstrating that this second binding site contributes essentially to the PI(4,5)P_2_-dependent membrane association of tubby. The positive charge cluster is conserved throughout TULP family proteins, indicating that simultaneous PI(4,5)P_2_ binding by two binding sites is a conserved feature of tubby family proteins that may be related to their cellular function. Thus, we hypothesize that cooperative PI(4,5)P_2_ binding may facilitate delivery of cargo into primary cilia by TULP proteins, which involves binding to the PI(4,5)P_2_-rich PM and subsequent dissociation from the PI(4,5)P_2_-depleted ciliary membrane.

## MATERIAL AND METHODS

### Molecular dynamics simulations

#### System setup and simulation details

All simulations were performed using the CG force field Martini 3 (open beta version (Souza and Marrink, 2020)) and the program package Gromacs (version 2018.1) (Abraham et al., 2015). We used the tubbyCT crystal structure (pdb code: 1I7E) (Santagata et al., 2001) and modeled the missing loops with the I-TASSER server (Yang et al., 2015). To generate the CG model, the missing loops were ligated to the crystal structure. This step was necessary to maintain the side chain orientations around the PI(4,5)P_2_ binding pocket of tubbyCT identified in the crystal structure. Based on the Martini protein model without any elastic network (de Jong et al., 2013), sidechain corrections (Herzog et al., 2016) were added. To maintain the secondary and tertiary protein structure, a Gō-like model was employed according to the procedure in reference (Poma et al., 2017). Based on contacts defined by an overlap and contacts of structural units-criterion evaluated only for the residues resolved in the crystal structure, Lennard-Jones interactions were added up to a cutoff distance of 1.1 nm (Thallmair et al., 2019). The dissociation energy of the Lennard-Jones potential was set to ε = 12.0 kJ/mol (Souza et al., 2019). In total, 541 Gō-like bonds were added for tubbyCT.

Different membrane compositions were used to study the PI(4,5)P_2_ binding of tubbyCT: (i) A simple POPC membrane containing one additional PI(4,5)P_2_ lipid which was embedded in the pocket identified in the crystal structure in the starting structure. (ii) In a second setup, the membrane consisted of POPC and PI(4,5)P_2_ lipids in a molar ratio of 95:5. (iii) As control setup, POPC membranes with 5 mol% of different negatively charged lipids (POPG, POPS, and PI(4)P) were used to test the impact of other negatively charged lipids. Also in case (ii) and (iii), one PI(4,5)P_2_ lipid was initially embedded in the crystal binding pocket of tubbyCT. In all simulations, the recently refined bonded parameters of the PI(4,5)P_2_ lipids were used (Sun et al., 2020). The membrane patches had a size of 15 × 15 nm^2^ containing approximately 700 lipids in total. They were generated using the program *insane.py* (Wassenaar et al., 2015). Finally, the system was neutralized and solvated in a 0.15 M NaCl solution. The rectangular box with an initial size of 15 × 15 × 14 nm^3^ contained ~17,300 water beads corresponding to ~69,200 water molecules.

The simulation parameters were chosen in accordance to the new reaction-field settings given in reference (de Jong et al., 2016). After the equilibration, setup (i) was simulated for 1 μs (10 replicas), setup (ii) for 5 μs (3 replicas), and setup (iii) for 1 μs in the case of POPG and POPS and for 5 μs in the case of PI(4)P (3 replicas for each lipid type).

For comparison, we also simulated a PH domain, which is a common protein binding PI lipids. Here, we employed the crystal structure of the PLCδ1-PH domain (pdb code: 1MAI) (Ferguson et al., 1995), a well-characterized stable PI(4,5)P_2_ binder (Stauffer et al., 1998; Várnai and Balla, 1998). We modeled the missing termini with the I-TASSER server (Yang et al., 2015) and ligated them to the crystal structure. The Gō-like model was set up similar to the tubbyCT resulting in 234 added Lennard-Jones interactions. Simulations were performed using the setup (i) and (ii) as described earlier for tubbyCT.

#### Analysis

To identify the unbinding of the proteins from the membrane, the distance between the plane of the phosphate groups of the lipids (PO4 beads) and the crystal binding pocket was calculated using the *gmx distance* tool. The phosphate plane is formed by the PO_4_^-^ groups which are directly linked to the glycerol moiety. The phosphate groups of the inositide head groups are not considered here. This procedure has the advantage that an exchange of individual PI(4,5)P_2_ lipids in contact with the protein does not alter the calculated distance.

To obtain more detailed information about the position of the PI(4,5)P_2_ lipid with respect to the protein surface, the distance between the PI(4,5)P_2_ head group and the crystal binding pocket was also calculated for the systems containing only one PI(4,5)P_2_ lipid (setup (i)). Again, the *gmx distance* tool was used.

To calculate the protein-PI(4,5)P_2_ contacts, the number of PI(4,5)P_2_ lipids with a distance ≤ 0.5 / 0.7 / 0.9 nm between the head group and the protein was analyzed (using *gmx select*) while the protein was considered being bound to the membrane. The number of contacts to the individual residues was also calculated using a distance cutoff of 0.5 nm.

#### Potential of mean force (PMF) calculations

To estimate the binding free energy of wildtype tubbyCT, tubbyCT R301A, and PLCδ1-PH domain, we calculated the PMF using umbrella sampling. We followed the procedure described in reference (Naughton et al., 2016). In brief, we generated conformations along the reaction coordinate by pulling the protein center of mass away from the PI(4,5)P_2_ head group perpendicular to the membrane plane (z-direction; force constant 1000 kJ/(mol nm^2^); pulling rate 0.001 nm/ps) while the PI(4,5)P_2_ lipid was restrained in space using a harmonic potential (force constant 1000 kJ/(mol nm^2^)) and the PI(4,5)P_2_ head group and the protein center of mass were kept at the same point in the plane of the membrane (x/y plane; force constant 100 kJ/(mol nm^2^)). For the umbrella sampling simulations, the harmonic constraint of the PI(4,5)P_2_ lipid was released and the distance between the PI(4,5)P_2_ head group and the protein center of mass was constrained with a harmonic potential (force constant 1000 kJ/(mol nm^2^)). This distance was sampled between 1.6–5 nm for tubbyCT and 1.2–4.6 nm for the PLCδ1-PH domain, respectively, using an interval of 0.1 nm. The resulting 35 windows were sampled for 2 μs in the case of tubbyCT and 1 μs in the case of the PH domain. The sampling for tubbyCT was increased to better distinguish the wildtype tubbyCT from its R301A mutant. To calculate the PMFs, the *gmx wham* tool was employed; error estimation was done using a bootstrap analysis with 100 bootstraps (Hub et al., 2010).

### Experimental assessment of membrane binding in living cells

#### Cell culture

CHO dhFr^-^ cells were cultured in MEM Alpha medium (gibco, ThermoFisher Scientific, Waltham, US) supplemented with 10% fetal calf serum, 1% penicillin and 1% streptomycin. Cells were seeded on glass bottom dishes and kept at 37°C and 5 % CO2. Two days after seeding, cells were transfected using JetPEI^®^ DNA transfection reagent (Polyplus Transfection, Illkirch-Graffenstaden, France) according to the manufacturer’s instructions. Experiments were performed 24h post-transfection.

#### Molecular biology

Expression constructs used for transfection were: mouse tubbyCT (AA 243-505) in pEGFP-C1 (NM_021885.4); Ci-VSP and Ci-VSP-C363S in pRFP-C1 (AB183035.1). Mutagenesis of of tubbyCT was done using QuikChange II XL Site-Directed mutagenesis kit (Stratagene, Agilent Technologies, Waldbronn, Germany).

#### Wide-field fluorescence microscopy

Experiments were performed on a Dmi8 upright microscope (Leica, Wetzlar, Germany). Images were acquired with an ORCA-Flash4.0 C13440-20C camera (Hamamatsu photonics, Hamamatsu, Japan) controlled by LAS X software (Leica). For determination of membrane localisation of GFP-tubbyCT constructs, CHO cells were co-transfected with a catalytically inactive Ci-VSP (RFP-Ci-VSP C363S) as membrane marker. To quantify membrane localization of tubbyCT, line profiles across the cells were derived, and ratios of membrane-localized GFP-tubbyCT fluorescence intensities (averaged from the two intersections with the PM, defined by Ci-VSP C363S RFP fluorescence peaks) and cytosolic fluorescence were calculated.

#### Combined TIRF microscopy and voltage-clamp experiments

TIRF imaging was done on a Dmi8 upright microscope (Leica, Wetzlar, Germany) equipped with an Infinity TIRF module (Leica), a HC PL APO 100x/1.47 OIL objective (Leica) and a widefield laser (Leica). GFP fluorescence was excited at 488 nm and imaged through a GFP-T (505-555 nm) emission filter (Leica). Images were acquired every 4 s with an ORCA-Flash4.0 C13440-20C camera (Hamamatsu photonics, Hamamatsu, Japan) controlled by LAS X software (Leica). Simultaneously, cells were whole-cell patch-clamped for control of Ci-VSP activity as described previously (Halaszovich et al., 2009; Leitner et al., 2019). Briefly, voltage clamp recordings were done with an EPC 10 amplifier controlled by PatchMaster software (HEKA Elektronik, Lambrecht, Germany). Patch pipettes were pulled from borosilicate glass (Sutter Instrument Company, Novato, CA, USA) and had an open pipette resistance of 2-3 MΩ after back-filling with intracellular solution containing (mM) 135 KCl, 2.41 CaCl_2_, 3.5 MgCl_2_, 5 HEPES, 5 EGTA, 2.5 Na_2_ATP, 0.1 Na_3_GTP, pH 7.3 (with KOH), 290-295 mOsm/kg. Series resistance (Rs) typically was below 6 MΩ. Cells where held at −60 mV and depolarized in a stairstep command. During these experiments, the experimental chamber was continuously fed with an extracellular solution (5.8 mM KCl, 144 mM NaCl, 0.9 mM MgCl_2_, 1.3 mM CaCl_2_, 0.7 mM NaH_2_PO_4_, 5.6 mM D-glucose, 10 mM HEPES, pH = 7.4).

Analysis and statistical evaluation of obtained imaging data was done with IGOR Pro (WaveMetrics, Lake Oswego, OR, USA). Data are displayed as mean ± SEM. For statistical comparison Student’s t tests was applied with an asterisk indicating statistical significance at p < 0.05.

## RESULTS

### PI(4,5)P_2_ binding affinity evaluated for a single lipid

To gain insights into the PI(4,5)P_2_ binding behavior of tubbyCT, we performed CG MD simulations of tubbyCT bound to a single PI(4,5)P_2_ embedded in a POPC bilayer using the Martini force field. Figure 1a illustrates the system setup. Figure 1b shows the time evolution of the distance between the PO_4_ plane of the binding leaflet and the previously characterized binding pocket (Santagata et al., 2001) for ten simulations of 1 μs each. It can be clearly seen that tubbyCT unbinds from the PI(4,5)P_2_ lipid in all cases within 250 ns. In addition, no stable rebinding event to the membrane is observed within the simulation time.

**Figure 1.**
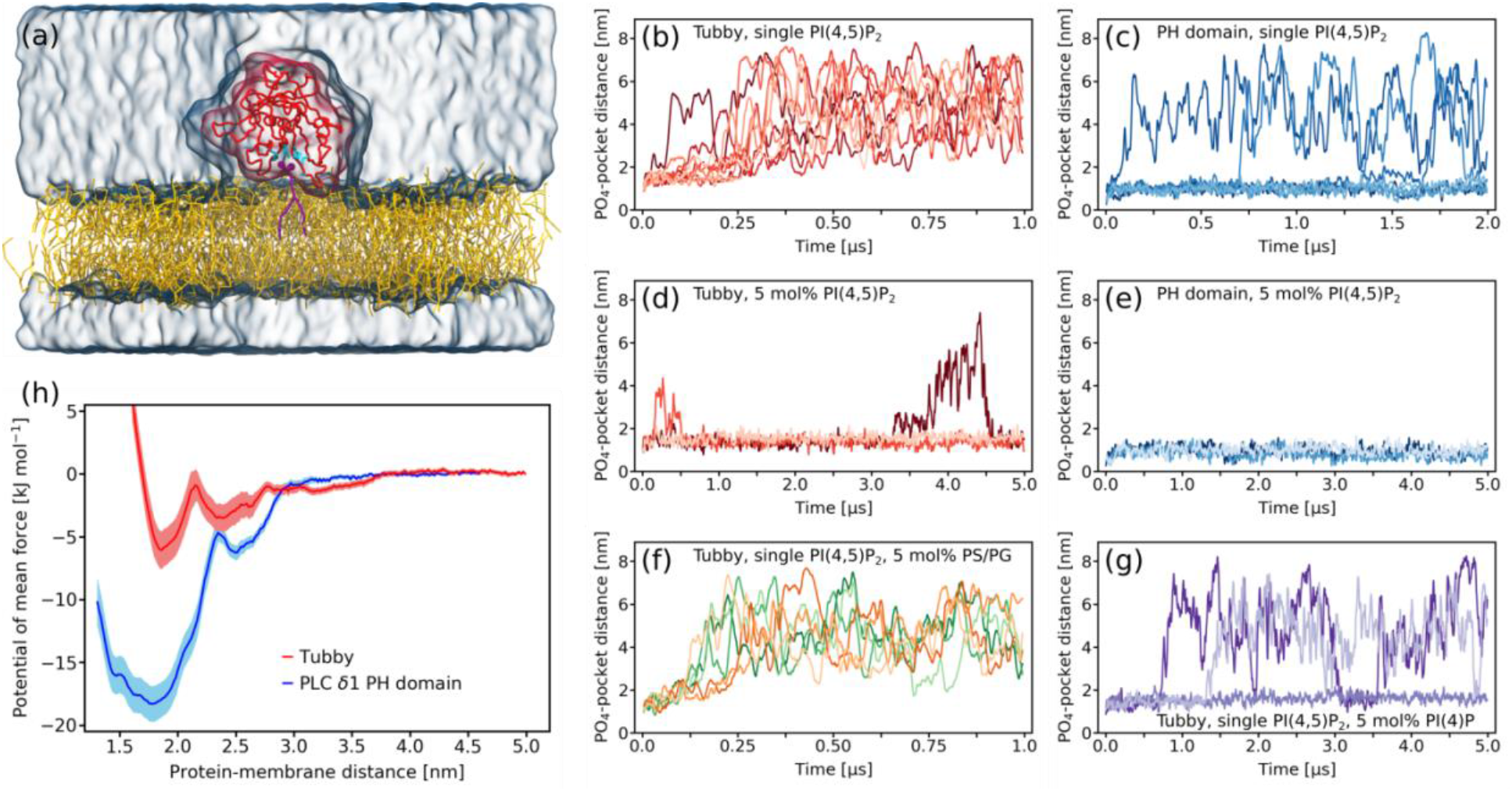
Binding a single PI(4,5)P_2_ lipid does not target tubbyCT stably to a model membrane. **(a)** CG system setup of tubbyCT (red) with one PI(4,5)P_2_ lipid (violet) in the binding pocket known from the crystal structure (cyan residues). The PI(4,5)P_2_ is embedded in a POPC bilayer (yellow); water and ions are shown as transparent surface. **(b,c)** Distance between the tubbyCT/PLCδ1-PH domain binding pocket and the phosphate layer (PO4 beads) of the binding leaflet containing a single PI(4,5)P_2_. Ten unbiased simulations of 1 μs each are shown. **(d,e)** Distance between the tubbyCT/PLCδ1-PH domain binding pocket and the phosphate layer (PO4 beads) of the binding leaflet containing 5 mol% PI(4,5)P_2_. Three unbiased simulations of 5 μs each are shown. **(f)** Control simulations of tubbyCT bound to one PI(4,5)P_2_ lipid embedded in a POPC membrane containing 5 mol% POPS (green) and POPG (orange) lipids, respectively. **(g)** Control simulations of tubbyCT bound to one PI(4,5)P_2_ lipid embedded in a POPC membrane containing 5 mol% PI(4)P. **(h)** Potential of mean force for the PI(4,5)P_2_ binding of tubbyCT (red) and PLCδ1-PH domain (blue).

As a reference for PI(4,5)P_2_ binding proteins, we also performed simulations of the PLCδ1-PH domain which is a well-known and stable PI(4,5)P_2_ binder (Stauffer et al., 1998; Várnai and Balla, 1998). An identical system setup like in the case of tubbyCT was used. Figure 1c shows the time evolution of the PO_4_-binding pocket distance for seven simulations of 2 μs each. The PLCδ1-PH domain binds much more stably to the PI(4,5)P_2_ lipid. Despite the doubled simulation time, unbinding was only observed in three cases and one rebinding event occurred.

To clarify whether an increased PI(4,5)P_2_ concentration impacts the binding stability of both proteins, we increased the PI(4,5)P_2_ concentration in the bilayer to 5 mol% and performed three simulations of 5 μs each. The observed time evolution of the PO_4_-binding pocket distance is shown in Figures 1d (tubbyCT) and 1e (PLCδ1-PH domain), respectively. Both PI(4,5)P_2_ sensors bind more stably to the membrane containing 5 mol% PI(4,5)P_2_. However, the increase in binding stability is more pronounced in the case of tubbyCT, where a single PI(4,5)P_2_ was not sufficient to target the protein for more than 250 ns to the bilayer surface. Moreover, observed unbinding events of tubbyCT from the membrane with high PI(4,5)P_2_ concentration are only transient and followed by rebinding.

We next asked, whether the increased membrane binding is specific for higher PI(4,5)P_2_ concentrations, or whether any negatively charged lipid could support the binding of tubbyCT to a single PI(4,5)P_2_. This is particularly relevant because the inner leaflet of the PM contains several negatively charged lipid species (Ingolfsson et al., 2014; Lorent et al., 2020). We decided to test three phospholipids abundant in the eukaryotic PM, POPS and POPG, as well as the singly phosphorylated phosphoinositide PI(4)P. The PI(4,5)P_2_ bound to tubbyCT was embedded in a POPC bilayer containing 5 mol% of either POPS, POPG, or PI(4)P. Figure 1f shows the PO_4_-binding pocket distance for three simulations of 1μs with POPS (green) and POPG (orange), respectively. In each case, no additional stabilization of tubbyCT membrane binding was observed. PI(4)P clearly increased membrane binding (Figure 1g), however, the stabilization was less pronounced than with the same concentration of PI(4,5)P_2_.

Our unbiased simulations discussed so far showed that the tubbyCT binds PI(4,5)P_2_ less strongly than the PLCδ1-PH domain. In order to get a quantitative comparison of the PI(4,5)P_2_ affinity of the two proteins, we calculated the potential of mean force (PMF) for the binding of the protein to the PI(4,5)P_2_ lipid. Figure 1h shows the resulting free energy profiles. They confirm that tubbyCT binds much weaker to PI(4,5)P_2_. The PMF minima differ by a factor of 3 (tubbyCT −6.1 kJ/mol; PLCδ1-PH domain −18.3 kJ/mol). The total binding free energy *ΔG*_bind_ can be calculated by integrating the PMF profile over the region of the bound state while taking into account the constraints in the membrane plane (Doudou et al., 2009). This yields a *ΔG*_bind_ = −2.4 kJ/mol for tubbyCT and *ΔG*_bind_ = −15.3 kJ/mol for the PLCδ1-PH domain. These findings confirm experimental data where PI(4,5)P_2_ affinity of tubbyCT and PLCδ1-PH was measured via gradual activation of a voltage-sensitive phosphatase (VSP) (Halaszovich et al., 2009; Leitner et al., 2019).

### Identification of a new PI(4,5)P_2_ biding site of tubbyCT

The CG MD simulations allow a microscopic investigation of the membrane binding behavior of tubbyCT and PLCδ1-PH domain. Figure 2a depicts a normalized histogram of the distance between the PI(4,5)P_2_ head group and the crystal structure binding pocket for the simulations with a single PI(4,5)P_2_ embedded in a POPC bilayer. Note that this distance is different from the distances depicted in Figure 1b–g where the distance between the PO_4_ plane of the binding leaflet and the binding pocket was analyzed. The distance to the head group shown here provides more detailed microscopic information about the protein-PI(4,5)P_2_ contacts.

**Figure 2.**
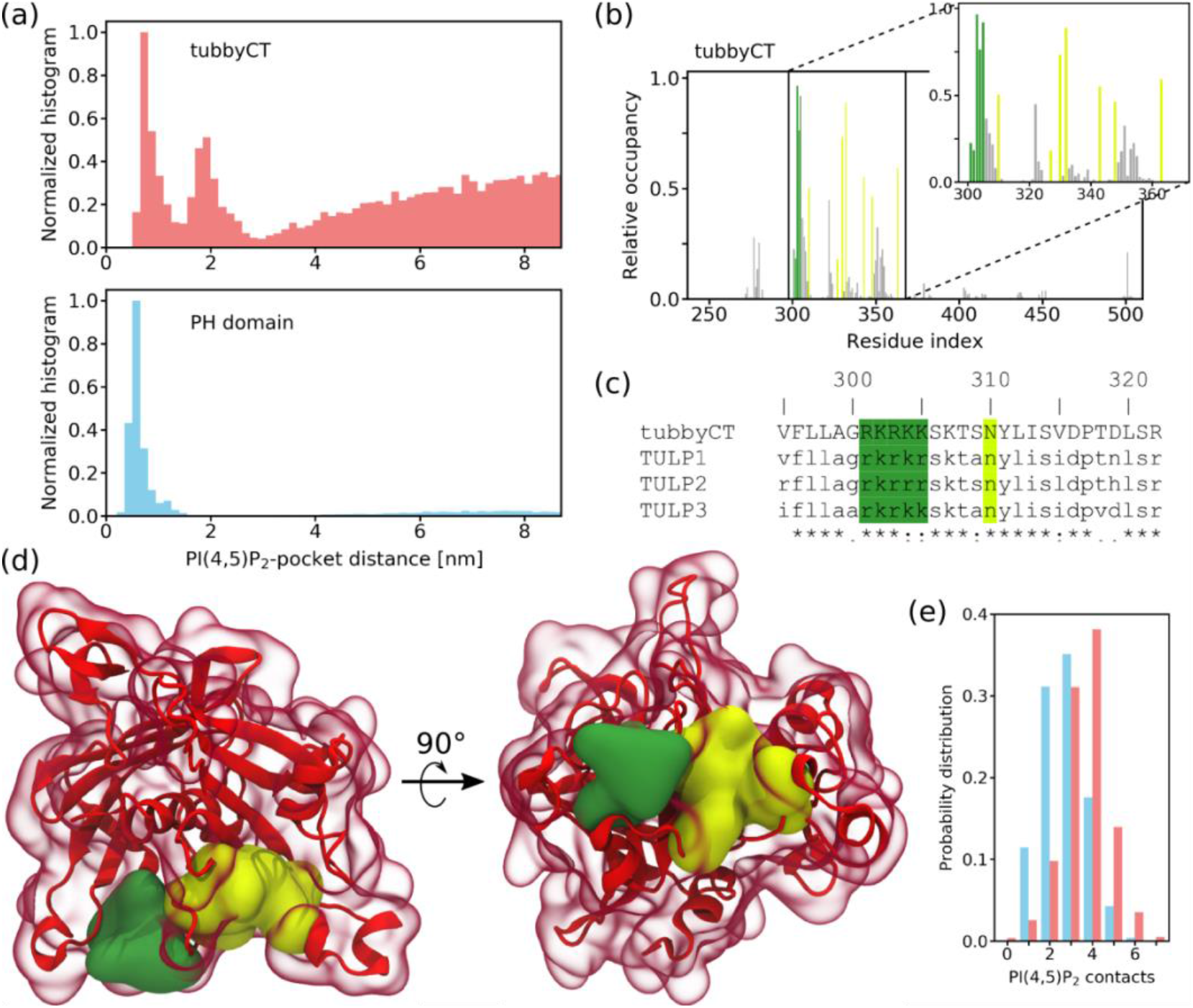
Identification of the second PI(4,5)P_2_ binding hot spot of tubbyCT. **(a)** Normalized distribution for the distance between the PI(4,5)P_2_ head group and the crystal structure binding pocket of tubbyCT (upper panel) and PLCδ1-PH domain (lower panel). The simulations were performed with one PI(4,5)P_2_ in the bilayer. **(b)** Relative PI(4,5)P_2_ occupancy of the tubbyCT residues calculated with a distance cutoff of 0.5 nm. Lemon bars represent residues of the crystal structure binding pocket; green bars highlight residues of the binding hot spot identified here. **(c)** Sequence alignment of tubbyCT with other proteins of the TULP family; color code according to (b). **(d)** Crystal structure of tubbyCT including the modelled loops. The surface of the crystal structure binding pocket is shown in lemon; the surface of the new binding hot spot in green. **(e)** Probability distribution of the total PI(4,5)P_2_-protein contacts for tubbyCT (red) and PLCδ1-PH domain (blue) binding to a membrane with 5 mol% PI(4,5)P_2_.

The difference in height of the distributions at long distance is due to the different PI(4,5)P_2_ binding affinity of tubbyCT (upper panel in Figure 2a) and PLC δ1 PH domain (lower panel). Strikingly, the histogram for tubbyCT clearly shows two distinct maxima at which a stabilizing interaction exists while in the case of the PLCδ1-PH domain only one maximum appears. The two maxima in the case of tubbyCT are in agreement with the PMF (red line in Figure 1h). Note that due to technical reasons the distance coordinate is again defined differently in the PMF calculation which causes the shift of the distance value.

The second maximum indicated an additional PI(4,5)P_2_ binding site on the tubbyCT surface. To evaluate this more closely, we calculated the average PI(4,5)P_2_ occupancy of each residue of tubbyCT for the simulations in which tubbyCT interacted with a membrane containing 5 mol% PI(4,5)P_2_ (Figure 2b). The lemon bars highlight the residues of the binding pocket previously identified in the crystal structure (Santagata et al., 2001). Besides those residues, there are five consecutive positively charged residues (AA 301–305, colored in green) of which in particular residues 303–305 exhibit a strikingly high PI(4,5)P_2_ occupancy. Accordingly, these residues constitute the second binding site identified in the PI(4,5)P_2_ head group-canonical binding site distance histogram (Figure 2a). To the best of our knowledge, this binding site has not been discussed so far. The sequence alignment of tubby-like proteins (TULPs) depicted in Figure 2c reveals that these five positively charged residues are conserved within the TULP family. The newly identified binding site is located in proximity to the crystal structure binding pocket and importantly, both PI(4,5)P_2_ binding sites are oriented in parallel so that simultaneous PI(4,5)P_2_ binding is possible (see Figure 2d). The surface of the two binding sites is colored according to Figure 2b. Because the binding sites are oriented adjacently on the same side of the protein surface, both of them can be occupied simultaneously when tubbyCT binds at a membrane with a sufficiently high PI(4,5)P_2_ concentration. Thus, the binding affinities of both binding sites act together and target tubbyCT to the membrane.

Figure 2e depicts the probability distribution of the number of simultaneous protein-PI(4,5)P_2_ contacts with a distance of ≤0.5 nm for tubbyCT (red) and the PLCδ1-PH domain (blue) when binding to a membrane with 5 mol% PI(4,5)P_2_. Multiple contacts to the same lipid are only counted once. It can be clearly seen that in the case of tubbyCT the distribution is shifted to a higher number of protein-PI(4,5)P_2_ contacts. The average number of PI(4,5)P_2_ lipids which are in contact with tubbyCT are listed Table 1 for cutoff distances of 0.5, 0.7, and 0.9 nm. The relative concentration of PI(4,5)P_2_ lipids which are in direct contact with tubbyCT (distance ≤0.5 nm) is 44%. It decreases to 19% for a distance of ≤0.9 nm which is still about four times higher than the PI(4,5)P_2_ concentration in the membrane (5%). For the simple model membrane used in the CG MD simulations here, the lipid fingerprint – i.e. the lipid composition of the annular lipid shell – of tubbyCT is highly enriched of PI(4,5)P_2_ while POPC is depleted. On average, the PLCδ1-PH domain is in contact with less PI(4,5)P_2_ lipids (see Table 1). However, the lipid fingerprint still shows an enrichment of PI(4,5)P_2_ in the vicinity of the PLCδ1-PH domain (34%).

**Table 1.** Number of lipids in contact with tubbyCT and the PLCδ1-PH domain depending on the cutoff distance.

| | tubbyCT | | | | PLCδ1-PH domain | | | |
| --- | --- | --- | --- | --- | --- | --- | --- | --- |
|  | PI(4,5)P<sub>2</sub> |  | POPC |  | PI(4,5)P<sub>2</sub> |  | POPC |  |
| Cutoff [nm] | abs. | rel. | abs. | rel. | abs. | rel. | abs. | rel. |
| 0.5 | 3.4±0.2 | 0.44 | 4.4±0.4 | 0.56 | 2.7±0.2 | 0.34 | 5.4±0.2 | 0.66 |
| 0.7 | 3.6±0.2 | 0.31 | 8.0±0.7 | 0.69 | 2.9±0.2 | 0.25 | 8.4±0.2 | 0.75 |
| 0.9 | 3.9±0.2 | 0.19 | 16.4±1.0 | 0.81 | 3.1±0.3 | 0.18 | 14.6±1.0 | 0.82 |
| Membrane composition | 18 | 0.05 | 334 | 0.95 | 18 | 0.05 | 334 | 0.95 |

### Characterization of the secondary binding site

To evaluate the role of the newly identified second binding site in membrane association of tubbyCT in living cells, we mutated the positively charged amino acids 301–305 that constitute the new binding site to alanine, which lacks electrostatic attraction to the anionic PI(4,5)P_2_. GFP-fused tubbyCT and the various mutants were expressed in CHO cells examined for their membrane localization by live-cell epifluorescence microscopy. As shown in Figure 3a and b, each single point mutant R301A, R303A, and K304A lost the strong membrane association of the wildtype domain and predominantly localized to the cytoplasm and nucleus. TubbyCT K305A showed a similar phenotype, although this mutant retained distinct, but reduced, membrane localization. We did not analyze position R302 by mutation as it binds D499 by hydrogen bonding and thus might be important for the tubbyCT secondary structure. Combinatorial neutralization of two (R301A/K304A or R303A/K305A), three (R303A/R304A/K305A) or four (R301A/R303A/R304A/K305A) of the site’s positive charges resulted in full localization to cytoplasm and nucleus and lack of detectable membrane association (Figure 3a, b).

**Figure 3.**
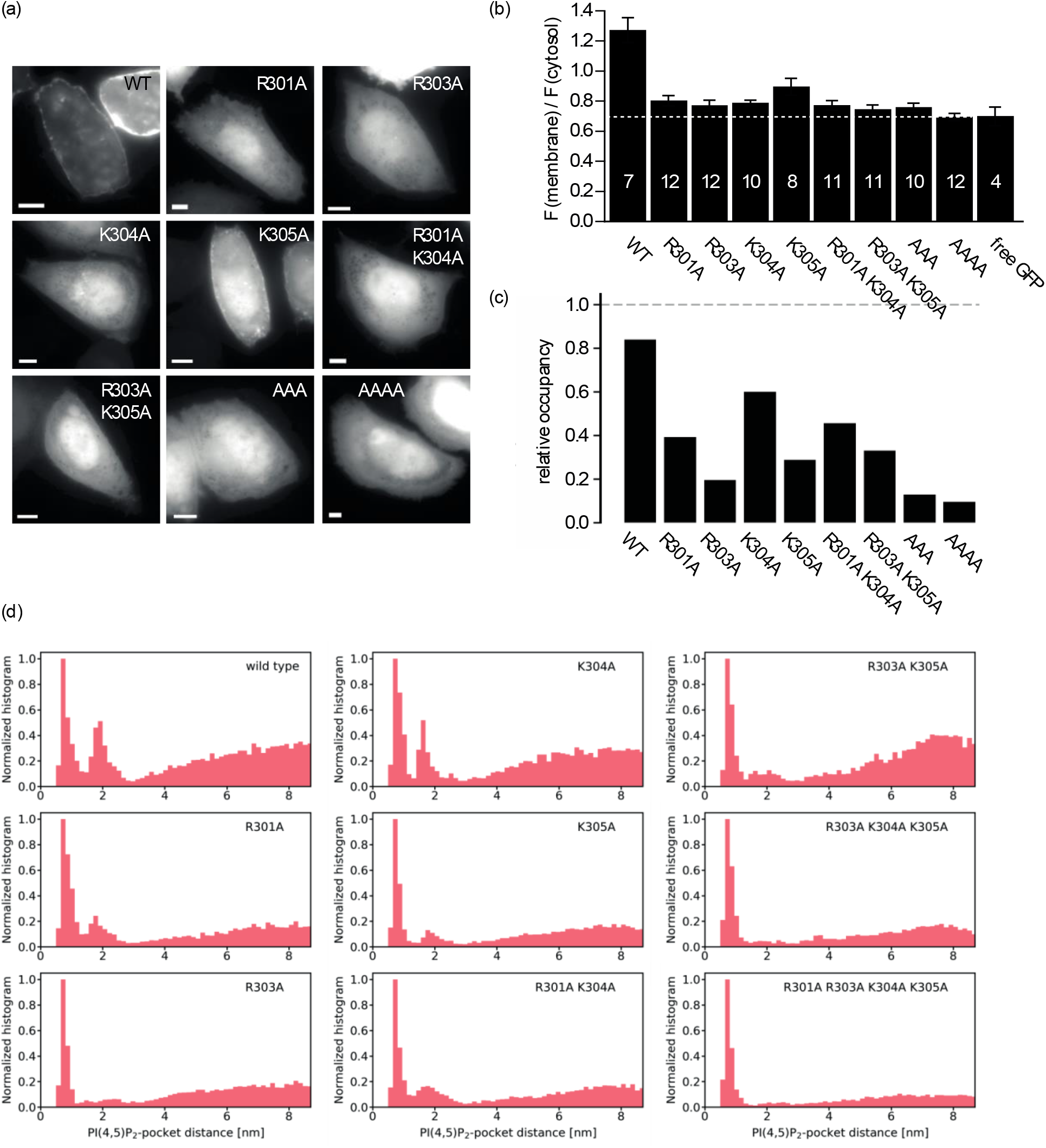
Mutational analysis of the second binding site. **(a)** Representative fluorescence images of tubbyCT wildtype and mutants expressed in CHO cells show the different degrees of membrane localization of the constructs (Scale bars, 5 μm). **(b)** Membrane-to-cytosol fluorescence ratios obtained from images as shown in (a). Localization of the membrane was defined as the local fluorescence maximum of a co-expressed RFP-fused membrane protein (Ci-VSP). **(c)** Ratio of occupancy of the second binding site relative to the canonical site obtained from occupancy distributions as shown in (d). **(d)** Normalized occupancy distribution for the distance between the PI(4,5)P_2_ head group and the canonical binding pocket of tubbyCT mutants. Simulations were performed with one PI(4,5)P_2_ in the bilayer.

In a complimentary manner, we investigated PI(4,5)P_2_ binding of the mutants by CG MD simulations. We simulated ten replicas of 1 μs simulation time for each tubbyCT mutant where the protein was initially bound to a single PI(4,5)P_2_ lipid embedded in a POPC bilayer. As a measure for PI(4,5)P_2_ binding behavior, we calculated the distance between the PI(4,5)P_2_ head group and the canonical binding site over the duration of each simulation. For tubbyCT WT, the histogram of the resulting distances shows two maxima corresponding to the two binding sites (c.f., Figure 2a). The occupancy of the second maximum was reduced by the various mutations (Fig. 4d). To estimate the relative binding strength of the second binding site in comparison to the canonical one, we integrated both maxima and calculated their relative occupancy, i.e. the ratio of the integrals of both binding site maxima. While for tubbyCT WT the relative occupancy is > 0.8, all mutants show a reduced relative occupancy with the triple and quadruple mutants exhibiting the lowest relative occupancy close to zero.

**Figure 4:**
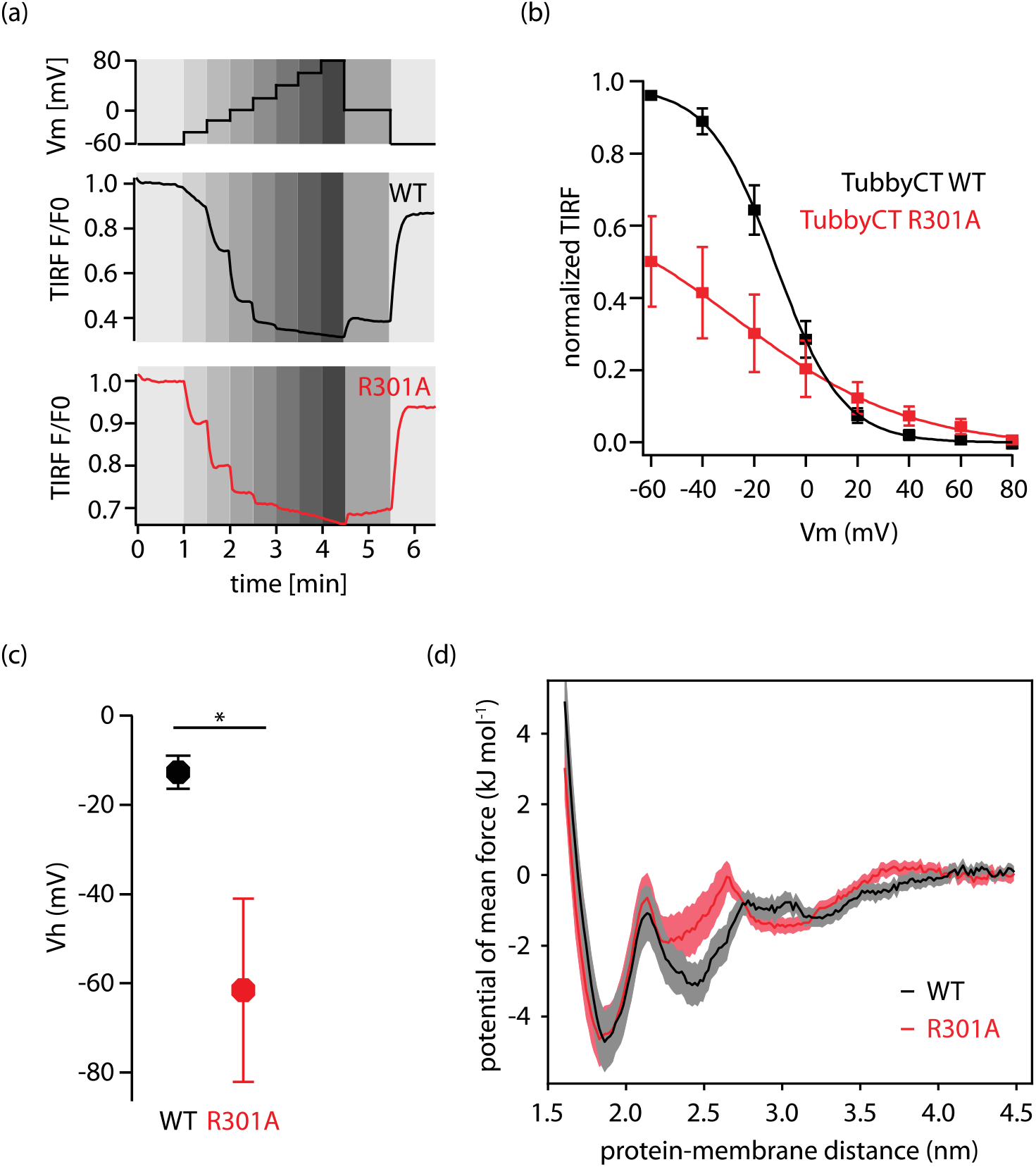
Contribution of the second binding site to PI(4,5)P_2_ affinity of tubbyCT. **(a)** Changes in membrane association of GFP-tubbyCT in response to activation of Ci-VSP as recorded by TIRF-M. CHO cells co-transfected with GFP-tubbyCT and RFP-Ci-VSP were whole-cell voltage-clamped and depolarized gradually (staircase voltage protocol, upper panel) while measuring membrane-localized fluorescence by TIRF-M. Lower panels show representative TIRF recordings for WT and R301A mutant, normalized to resting signal at −60 mV. **(b)** Fluorescence-voltage curves obtained from experiments as in (a) were fitted by a Boltzmann function and normalized to maximal fitted fluorescence change. Shown are averaged data from N = 9 and 7 cells for WT (black) and R301A (red) respectively. **(c)** Mean voltage required for half-maximal dissociation from the membrane from curves shown in (b). **(d)** Potential of mean force for the PI(4,5)P_2_ binding of tubbyCT WT (black) and tubbyCT R301A (red).

Next, we chose tubbyCT R301A as a representative mutant for a more detailed experimental characterization of the PI(4,5)P_2_ binding affinity in living cells. We used the voltage-sensitive phosphatase (from *Ciona intestinalis;* Ci-VSP) for gradual and stepwise change of the PM level of PI(4,5)P_2_ in living CHO cells. VSPs are 5-phosphatases that dephosphorylate PI(4,5)P_2_ to PI(4)P with a gradual dependency of their enzymatic activity on the membrane potential, where depolarization increases the activity. As the PI(4,5)P_2_ concentration at each imposed membrane voltage depends on the counter-acting activities of the VSP and intrinsic PI4P-5-kinases, the step-wise activation of Ci-VSP (Figure 4a, upper panel) allowed for titration of PI(4,5)P_2_ levels (cf., Halaszovich et al. 2009; Costa et al, 2015; Leitner et al., 2019). To this end, we co-expressed the respective GFP-tubbyCT mutant or wildtype together with RFP-tagged Ci-VSP. RFP-positive cells were whole-cell voltage-clamped while membrane association of GFP-tubbyCT constructs was monitored by TIRF microscopy. As shown in Figure 4a, incremental depolarization from −60 mV to +80 mV induced progressive dissociation of tubbyCT WT and R301A mutant from the PM, as reported by decreasing TIRF signal, consistent with the membrane binding being PI(4,5)P_2_-depedendent. TubbyCT R301A translocation occurred at lower membrane potentials than translocation of tubbyCT WT, indicating reduced PI(4,5)P_2_ affinity (Figure 4a,b). Of note, the lower overall reduction in normalized signal amplitude observed with the mutant is also consistent with the lower basal membrane association (Figure 3a). For quantitative assessment, we fitted TIRF signal amplitudes with a Boltzmann function that describes the voltage dependency of VSP enzymatic activity (e.g., Halaszovich et al., 2009). As shown in Figure 4b,c, half-maximum translocation of the R301A mutant occurred at much more negative potentials (V_h_ = −61.5 ± 20.6 mV) compared to the WT construct (V_h_ = −12.7 ± 3.7 mV). Thus, much less activation of the phosphatase was required for unbinding of the mutant domain, which is equivalent to the dissociation at a higher PI(4,5)P_2_ concentration.

Finally, we quantified the impact of the R301A mutation on the PI(4,5)P_2_ binding free energy, by calculating the PMF of the R301A mutant and WT tubbyCT. Figure 4d shows that the potential depth at the second binding site (protein-membrane distance of 2.4 nm) is reduced while the canonical binding site (at 1.8 nm) is unaffected. The total binding free energy *ΔG*_bind_ calculated from the PMF profile yields a *ΔG*_bind_ = −2.4 kJ/mol for tubbyCT WT and *ΔG*_bind_ = −2.0 kJ/mol for the R301A mutant. Despite the small difference, it confirms the reduced affinity measured in living CHO cells.

In summary, experimental and computational analyses agree in showing that the PI(4,5)P_2_ affinity of tubbyCT critically depends on the novel second binding site, such that PM association at physiological PI(4,5)P_2_ concentration requires lipid interaction at both binding sites.

## DISCUSSION

Here, we identify a second PI(4,5)P_2_-binding site in the C-terminal domain (‘tubby domain’) of the tubby protein. It consists of a conserved cluster of positively charged amino acids and is located next to the classical, or canonical binding site at the relatively planar protein-lipid interface of the tubby domain. PI(4,5)P_2_ binding by the canonical binding site as previously shown by crystallography as well as by mutational analysis (Santagata et al., 2001) was fully reproduced by our CG MD simulations. Yet, occupancy of the second binding was nearly as high (ratio 0.8) as the occupancy of that primary site, and indeed proved essential for PI(4,5)P_2_ binding and hence membrane association of the domain at physiological PI(4,5)P_2_ levels.

### PI(4,5)P_2_ binding mode of the tubby domain

Interestingly, polybasic motifs similar to the second binding site mediate membrane association of many cytosolic proteins by electrostatic interactions with PI(4,5)P_2_ and PIP_3_ (Heo et al., 2006; Sun et al., 2018) or PI(4)P (Hammond et al., 2012). Here we observed substantial specificity of the new binding site for PI(4,5)P_2_. Thus, PI(4)P had a much weaker, although detectable capacity to bind tubbyCT to the membrane, and the anionic PS and PG were ineffective. Such specificity may help to ensure targeting to the PM rather than to other negatively charged membrane compartments.

How does the dual binding mode of tubbyCT compare to well-characterized PI-binding domains? PH domains bind their phosphoinositide ligand by a single binding pocket (Ferguson et al., 1995). The high selectivity for distinct PI species characteristic for some PH domains is achieved by stereospecificity of the interaction between the binding site and the anionic headgroup of the ligand and differences in binding selectivity arise from well-defined variations in the amino-acid sequence of the binding site (Ferguson et al., 2000; Ferguson et al., 1995; Lietzke et al., 2000; Park et al., 2008). However, the steric interaction with the anionic headgroup of the lipid also goes along with high-affinity binding of the isolated headgroup, e.g. the soluble second messenger IP_3_ in the case of the PI(4,5)P_2_-binding PH domain of PLCδ1 (Ferguson et al., 1995). This second cellular high affinity ligand is a major confounding problem in the use this domain as a reliable PI(4,5)P_2_ biosensor (Hammond and Balla, 2015; Hirose et al., 1999). In contrast, the requirement of the second binding site interaction at the membrane may contribute to the fact that tubbyCT has no detectable IP_3_ affinity (Quinn et al., 2008; Szentpetery et al., 2009).

Binding of the GRP1-PH domain to its specific ligand, PIP_3_, in the membrane is substantially enhanced by the presence of bulk anionic lipids such as PS and PtdIns. However, this interaction is thought to be mediated by a weak electrostatic interaction with the canonical binding site that precedes the high affinity binding of the specific substrate in a kind of ‘search mode’, rather than by a secondary interaction site (Corbin et al., 2004).

More similar to the two-binding-site mode identified here for tubbyCT, some PH domain proteins (Grp1, ARNO) feature a polybasic motif outside of the PH domain which enhances affinity for negatively charged membranes in a cooperative manner (Nagel et al., 1998; Santy et al., 1999). Some other classes of phosphoinositide recognition domains, such as the FYVE and PX domains, also use a dual binding mode where in addition to headgroup recognition insertion of a hydrophobic moiety into the membrane mediates membrane association and increases affinity for the PI ligand in the membrane environment (reviewed in (Lemmon, 2008)).

### Functional implications of the binding mode

The simultaneous binding of two PI(4,5)P_2_ molecules not only increases the overall affinity as shown here by mutations in one of the binding sites, but also predicts a steeper concentration dependence of membrane binding compared to binding by a single binding pocket as in the classical PH domains. Such a steep concentration dependence may be particularly relevant for differential membrane association under conditions of spatial or temporal PI(4,5)P_2_ inhomogeneities. E.g., tubby-domain-containing proteins may robustly bind at basal PM levels of PI(4,5)P_2_, while readily dissociating from the membrane at moderately decreased levels. Vice versa, cooperative binding may strongly favor binding to PI(4,5)P_2_-enriched membrane domains as opposed to the bulk membrane.

Given the conservation of the new binding site motif in mammalian TULP proteins (Fig. 2c), we consider this idea in the context of known common functions of TULP proteins. Although the cell biology of tubby-like proteins is only beginning to be illuminated, a congruent theme is the trafficking of proteins into cilia (Mukhopadhyay and Jackson, 2011). The following mechanistic model emerged (Badgandi et al., 2017): tubby domains can bind to cargo proteins designated for ciliary delivery by interacting with a ciliary localization signal of such proteins and this interaction requires membrane association of the tubby domain through PI(4,5)P_2_ binding (Badgandi et al., 2017). The TULP also binds to the intraflagellar transport complex (IFT-A), which shuttles the entire complex into the cilium (Mukhopadhyay et al., 2010). Once localized in the cilium, the cargo would be released from the TULP, because the ciliary membrane is poor in PI(4,5)P_2_ (Badgandi et al., 2017). The discrimination between different PI(4,5)P_2_ concentrations underlying this cyclic process may be facilitated by PI(4,5)P_2_ binding by two complementary sites.

## Author Contributions

Conceptualization, V.T., S.T., and D.O.; Investigation, V.T., L.S., and S.T.; Writing, S.T., D.O., and V.T.; Discussion: all authors; Funding Acquisition, D.O. and S.J.M.

## Acknowledgements

We like to thank Drs. Lawrence Shapiro and Yasushi Okamura for providing expression plasmids. S.T. thanks the Center for Information Technology of the University of Groningen for providing access to the Peregrine high performance computing cluster.

